# Molecular Evolution and Interaction of 14-3-3 Proteins with H^+^-ATPases in Plant Abiotic Stresses

**DOI:** 10.1101/2023.05.18.541295

**Authors:** Wei Jiang, Jing He, Mohammad Babla, Ting Wu, Tao Tong, Adeel Riaz, Fanrong Zeng, Yuan Qin, Guang Chen, Fenglin Deng, Zhong-Hua Chen

## Abstract

Environmental stresses severely affect plant growth and crop productivity. Regulated by 14-3-3 proteins (14-3-3s), H^+^-ATPases (AHA) are important proton pumps that can induce diverse secondary transport via channels and co-transporters for the abiotic stress response of plants. Many studies demonstrated the roles of 14-3-3s and AHAs in coordinating the processes of plant growth, phytohormone signaling, and stress responses. However, the molecular evolution of 14-3-3s and AHAs has not been summarized in parallel with insights across multiple plant species. Here, we review the roles of 14-3-3s and AHAs in cell signaling to enhance plant responses to diverse environmental stresses. We analyzed the molecular evolution of key proteins that are associated with 14-3-3s and AHAs in plant growth and hormone signaling. The results revealed evolution, duplication, contraction, and expansion of 14-3-3s and AHAs in green plants. We also discussed the stress-specific expression of those *14-3-3s* and *AHAs* in a eudicot (*Arabidopsis thaliana*), a monocot (*Hordeum vulgare*) and a moss (*Physcomitrium patens*) under abiotic stresses. We propose that 14-3-3s and H^+^-ATPases respond to abiotic stresses through many important targets and signaling components of phytohormones, which could be promising to improve plant tolerance to single or multiple environmental stresses.

**Highlight:** We review the response and adaptation of 14-3-3s and AHAs to diverse environmental stimuli and we analyze the evolutionary features and molecular functions of 14-3-3s and AHAs.

## Introduction

In recent years, increasing impact of climate change adversely affects natural and agricultural ecosystems in the world (Waadt et al., 2022). It results in higher frequency and severity of abiotic stresses including heat waves (Ding et al., 2020), cold (Gong et al., 2020), drought (Gupta et al., 2020), waterlogging (Khan et al., 2020), salinity (Chen et al., 2021c), and toxic metals (Zhang et al., 2022a). These abiotic stresses affect the global utilization of arable lands and severely limit crop productivity (Li et al., 2023). Plants have developed varied strategies to perceive and acclimatize to abiotic stresses during terrestrialization (Waadt et al., 2022; Zhao et al., 2019). Thus, understanding how plants sense stress signals and adapt to multiple environmental constraints conditions during the evolutionary history of 450 Ma of green plants is essential for sustaining global food security in the future (Chen, 2022; Jiang et al., 2022; Zhang et al., 2022a).

14-3-3 proteins (14-3-3s) were first discovered in bovine brain nearly 60 years ago (Moore & Perez, 1967). In plants, 14-3-3s are also named as G-box factor 14-3-3s (GF14s), general regulatory factors (GRFs), and Tomato Fourteen-Three-three (TFTs) (Zhao et al., 2021). 14-3-3s are 25-30 kD acidic proteins expressed in different tissues and organisms (Huang et al., 2022b). 14-3-3s play important role in modulating membrane transport [e.g., calcium (Ca^2+^) channels, inward rectifying K^+^ (K_in_) channels (KAT), H^+^-ATPases, and guard cell outward rectifying K^+^ (GORK)] (Cotelle& Leonhardt, 2015), hormone signaling (Camoni et al., 2018), nutrient and carbon metabolism (Shin et al., 2011; Zhao et al., 2021), and stomatal movements (Cotelle & Leonhardt, 2015) and modification of binding partners, including receptors and protein kinases (Cotelle & Leonhardt, 2015; Zhao et al., 2021). Environmental stresses affect the function of 14-3-3s in many ways (Denison et al., 2011): by activating signaling pathways and phosphorylation of corresponding proteins (Paul et al., 2012), influencing the activity of 14-3-3s and the signaling molecules [e.g., divalent cations and adenosine monophosphate (AMP)] (Jaspert et al., 2011), and altering the post-translational modification sites (PTMs) of 14-3-3 isoforms (de Boer et al., 2013; Wilson et al., 2016).

Autoinhibited H^+^-ATPases (AHAs) are adenosine triphosphate (ATP)-driven proton pumps that connect ATP hydrolysis with electrical proton gradients across cell membranes (Falhof et al., 2016; Fuglsang & Palmgren, 2021; Li et al., 2022b). AHAs contain five cytosolic domains, including N-terminal, actuator domain, phosphorylation domain, nucleotide binding domain, and regulation (R) domain (C-terminal autoinhibitory domain) (Li et al., 2022a). The N-terminal and R domain play roles in activity regulation, while the other domains are responsible for the catalytic activity (Miao et al., 2022). The *Arabidopsis thaliana* genome includes 11 AHA members (AHA1∼AHA11) that are localized in the plasma membrane (PM) except the tonoplast-localized AHA10 (Seidel, 2022). PM H^+^-ATPases consume the energy stored in ATP to shuttle protons to apoplast, which implicate a phosphorylation intermediate that form the conserved Asp residue in catalytic domain (Li et al., 2022b).

14-3-3 proteins, H^+^-ATPases, and their interactions play significant roles in response to many biotic (Li et al., 2022b; Ormancey et al., 2017) and abiotic (Falhof et al., 2016; Huang et al., 2022b) stresses (Fuglsang & Palmgren, 2021; Li et al., 2022a). For instance, in *A. thaliana*, 14-3-3κ/λ and salt overly sensitive 3 (SOS3) proteins selectively regulating SOS2 and SOS2-like protein kinase 5 (PKS5), which regulate Na^+^ homeostasis through coordinately mediating PM Na^+^/H^+^ antiporter (NHX) and AHA under salt stress (Yang et al., 2019). When sugar beet (*Beta vulgaris*) cells are exposed to cold stress, 14-3-3 proteins accumulate in the PM with increased H^+^-ATPase activities (Chelysheva et al., 1999). *Solanum lycopersicum* TFT4 plays a crucial role in alkaline stress through modulating H^+^-ATPase, auxin transport, and PKS5-DNAJ HOMOLOG3 pathway in root tips (Xu et al., 2013). Additionally, overexpression of maize (*Zea mays*) *ZmGF14-6* in rice confers drought tolerance via the strongly induced expression of genes associated with drought resistance (Campo et al., 2012). Overexpression of wheat (*Triticum aestivum*) *TaGF14b* (Zhang et al., 2018) and apple (*Malus domestica*) *MdGRF11* (Ren et al., 2019) in tobacco can enhance salt and drought tolerance. Moreover, most *Cucumis sativus CsGF14s* (e.g., *CsGF14a/b/d/e/f/g*) were up-regulated during drought and cold stress (Xu et al., 2021).

The molecular functions and evolution of 14-3-3s and H^+^-ATPases have been well summarized (Huang et al., 2022b; Jiang et al., 2023b; Li et al., 2022a; Li et al., 2022b; Ormancey et al., 2017). However, the evolution of 14-3-3s and AHAs have not been entirely explored in terms of environmental stresses. Here, we review the response and adaptation of 14-3-3s and AHAs to diverse environmental stimuli and we analyze the evolutionary features and molecular functions of 14-3-3s and AHAs. Furthermore, we discuss the interaction of 14-3-3s and AHAs with various targets and phytohormone signals simultaneously.

### Co-evolution of AHA and 14-3-3 proteins

#### Phylogeny, evolution, structure, and interaction of 14-3-3 proteins in plants

14-3-3 proteins are highly conserved among green plants might have originated from Chromista (e.g., *Proteomonas sulcata*), implying that the 14-3-3s in different organisms may share similar functions (Jiang et al., 2023b). However, the evolution and biological functions of the 14-3-3s in fern and moss under abiotic stress remain unclear. Herein, phylogeny, and motif alignment of 14-3-3s from land plants were performed to identify protein sequences in *Physcomitrium patens*, *Ceratopteris richardii*, *Oryza sativa*, *Hordeum vulgare*, and *A. thaliana* (Figs. 1A, B, S1). The 50 14-3-3 proteins could be divided into the epsilon and non-epsilon group (Fig. 1A). Nearly half fern and moss 14-3-3 members are in the epsilon group (Tian et al., 2015). Besides, most barley and rice 14-3-3s were located in the non-epsilon group (Ren et al., 2023; Wang et al., 2023c), and only one barley (Hv14-3-3F) (Mikhaylova et al., 2021) and two rice (OsGF14H/G) (Tian et al., 2015; Wang et al., 2023c) isoform were located in epsilon group (Fig. 1A). Interestingly, *C. richardii* harbors 12 14-3-3 proteins (Marchant et al., 2022), which might be associated with the genome expansion during evolution. Annexin-like motifs are highly conserved in 14-3-3 proteins in *P. patens*, *C. richardii*, *O. sativa*, *H. vulgare*, and *A. thaliana* (Fig. 1B), which might be pivotal for membrane phospholipid binding and Ca^2+^ binding (Pandey et al., 2021).

**Fig. 1.**
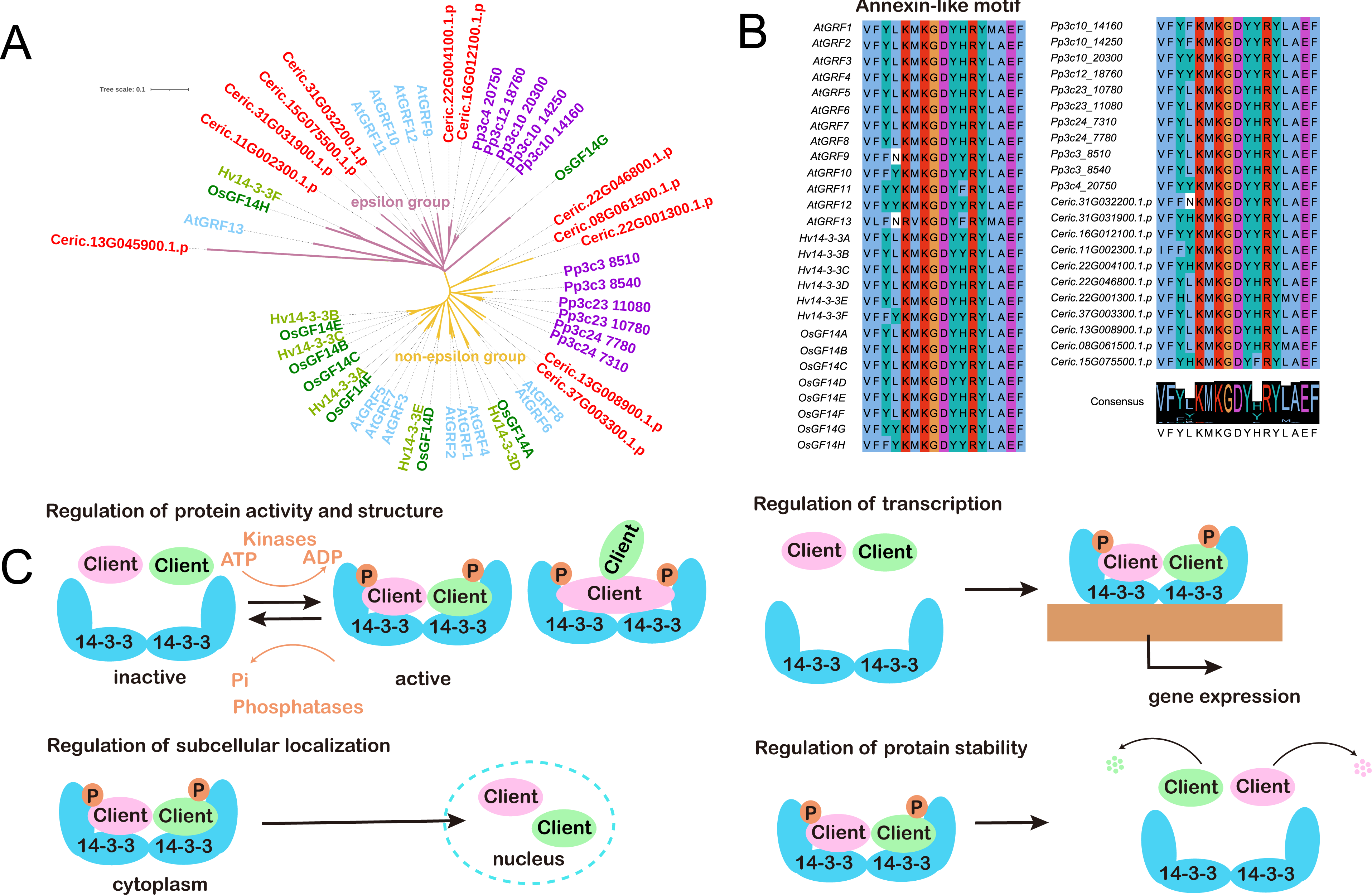
Molecular evolution of 14-3-3 proteins in plants. Phylogeny of 14-3-3 proteins in *Physcomitrium patens*, *Ceratopteris richardii*, *Oryza sativa*, *Hordeum vulgare*, and *Arabidopsis thaliana* (A). Motif alignment of 14-3-3 proteins in *Physcomitrium patens*, *Ceratopteris richardii*, *Oryza sativa*, *Hordeum vulgare*, and *Arabidopsis thaliana* (B). Schematic diagram of the structure and molecular interactions of 14-3-3 proteins (C).

The 14-3-3 proteins might regulate the structure (Fig. 1C), activity (Wilson et al., 2016) (Fig. 2), subcellular localization (Huang et al., 2022b), translocation (Fan et al., 2023; Liu et al., 2017), and stability of the client proteins (Camoni et al., 2018). In the 53 examined species of Chlorophyta and Embryophyta, the percentage of tandem, block, and duplication within this gene family were 8%, 39%, and 7%, respectively (Fig. 3) (https://bioinformatics.psb.ugent.be/plaza/versions/plaza_v5_monocots/genes/gene_d uplication_analysis/0/gf/HOM05M000404). 14-3-3s might undergo expansion in ferns (e.g., *C. richardii*) (Fig. 1) and mosses (e.g., *P. patens*) (Jiang et al., 2023b), while no duplication of 14-3-3s were found in monocots such as *Sorghum bicolor*, *H. vulgare*, and *Aegilops tauschii* (Fig. 3). The gene loss and duplication might be accompanied with the functional plasticity of 14-3-3s (Mikhaylova et al., 2021).

**Fig. 2.**
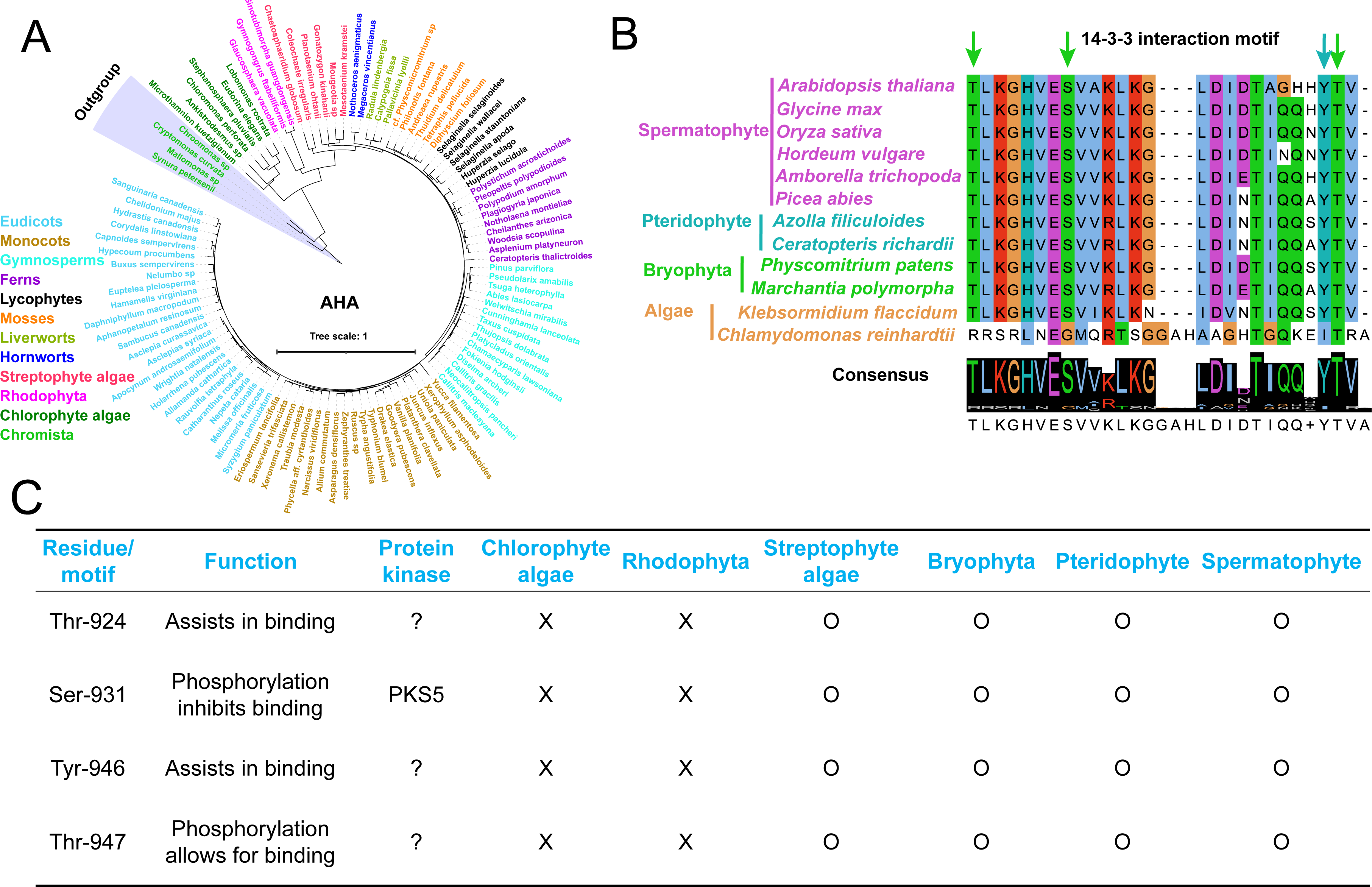
Evolutionary analysis of AHA proteins in land plants and algal species. The phylogenetic tree includes species in major clades in eudicots, monocots, gymnosperms, ferns, lycophytes, mosses, liverworts, hornworts, streptophyte algae, Rhodophyta, chlorophyte algae and Chromista (outgroup) (A). Motif alignment of AHA proteins in algae, Bryophyta, pteridophyte and spermatophyte (B). The interaction motif between AHA and 14-3-3 in green plants (C); motif according to Arabidopsis AHA2 nomenclature; ?, the protein kinase is not known; O, the proteins have this motif; ×, no this motif.

**Fig. 3.**
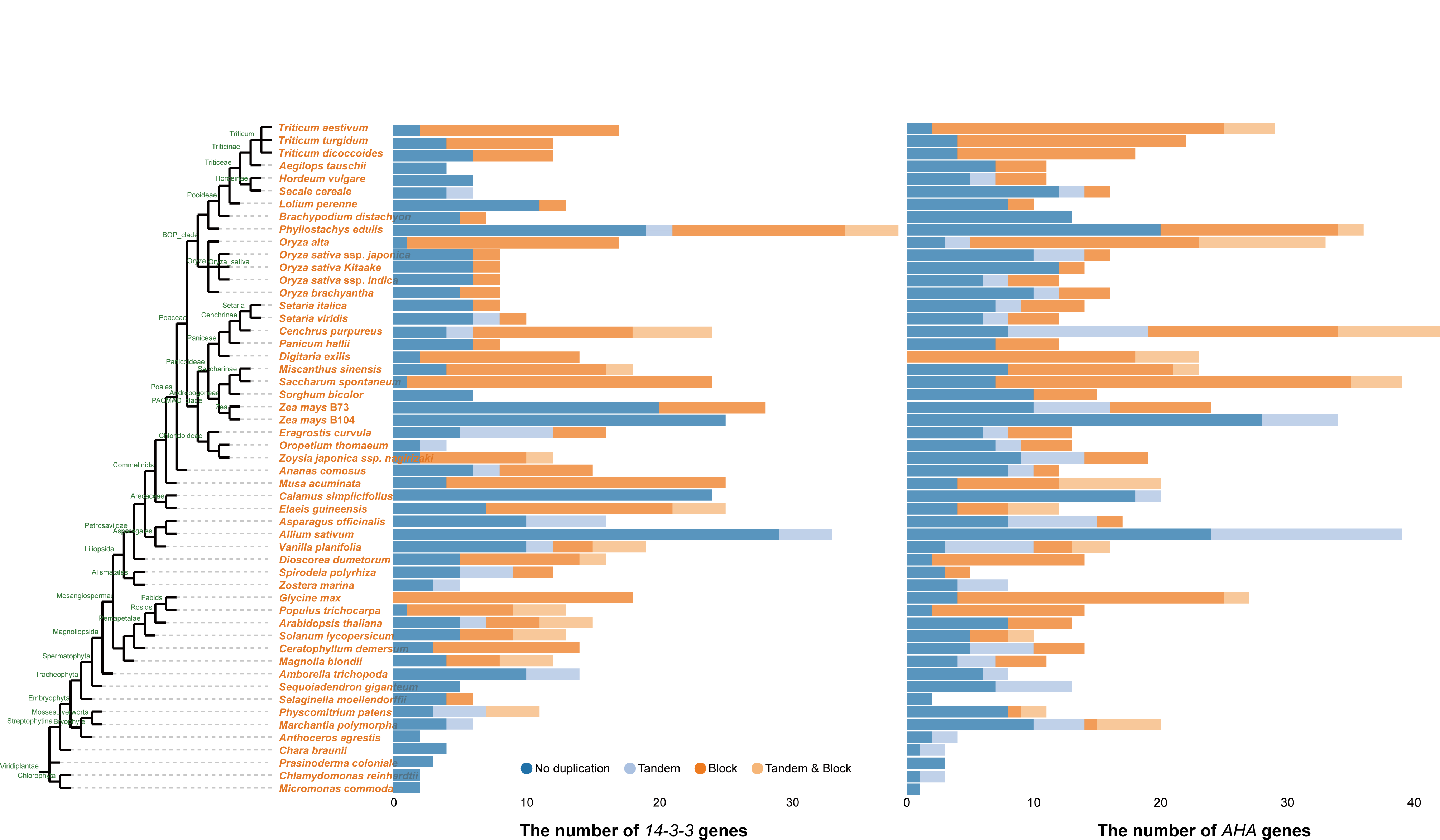
Tandem and block gene duplicate of *14-3-3s* and *AHAs* gene family in Chlorophyta and Embryophyta.

The 14-3-3s play essential role as “adapter protein” to regulated the interactions between client proteins, such as phosphatases, kinases, and signaling proteins (Shin et al., 2011). The interaction of 14-3-3s and AHAs have been studied in many plants (Li et al., 2022a) (Fig. 2). Proteomic analysis exhibited that the targets of 14-3-3s are related to ion transport (K^+^, Ca^2+^, Cl^-^) (Cotelle & Leonhardt, 2015) and phytohormone signaling [e.g. cytokinin, gibberellin (GA), ethylene (ETH), auxin (AUX), abscisic acid (ABA) and brassinosteroids (BR)] (Camoni et al., 2018; Huang et al., 2022b; Jaspert et al., 2011). Essential motifs of 14-3-3 proteins for interaction are protein kinase C pseudosubstrate, annexin-like, internal binding space, phosphorylation site, EF-hand (Helix E and Helix F calcium-binding) and nuclear export signal motif (Jiang et al., 2023b) (Fig. S1). The > 80% amino acid sequence similarity of 14-3-3s showed that those proteins are highly conserved in land plants with a slightly variable N/C-terminal and five conserved motifs that (Huang et al., 2022b; Mikhaylova et al., 2021) (Fig. S1). The conserved domains of the 14-3-3s displays similar function and the variable residues at the surface of the 14-3-3s show isoform specific function (Huang et al., 2022b).

#### Evolution of AHA and the interaction with 14-3-3 proteins

We found 860 *AHAs* in 53 Chlorophyta and Embryophyta species (https://bioinformatics.psb.ugent.be/plaza/versions/plaza_v5_monocots/gene_families/view/HOM05M000292), the percentage of tandem, block, and both gene duplicate in *AHA* family were 14%, 37%, and 8%, respectively (Fig. 3). The orthologs of AHAs, were confirmed in mostly green plants with high similarity to those in *A. thaliana* (Fig. S2). *Pinus taeda*, a gymnosperm species, had three isoforms of AHA proteins (Fig. S3), which may be associated with no whole-genome duplications (WGDs) in its evolutionary history. Besides, the 20, 26, 23, and 25 members of AHAs, respectively in *Marchantia polymorpha* (liverwort), *Phoenix dactylifera* (monocot), *Chenopodium quinoa* (eudicot) and *Glycine max* (eudicot) indicated that they might have experienced WGDs (Fig. S3). The expansion of AHA family in vascular plants (tracheophytes) might be due to the WGD events and polyploidy during evolution, which may be essential for adapting to changing climate (Li et al., 2022b). The putative AHA orthologues exist in land plants and algae (Fig. 2A). The origin of plants *AHA* gene family might be traced to Chromista (e.g., *Synura petersenii*, *Mallomonas sp*, and *Chroomonas sp*) (Fig. 2A), implying that AHAs are highly conserved in both green algae and plants.

In *A. thaliana*, 14-3-3s binds to the R domain of H^+^-ATPase at the C-terminal, affecting the H^+^-ATPase activity (Fuglsang & Palmgren, 2021; Steger et al., 2022). Additionally, the ε isoforms of 14-3-3 proteins (14-3-3ε/μ/ο) exhibit a lower affinity for binding with H^+^-ATPase than the non-ε 14-3-3 isoforms (14-3-3χ/κ/λ/ω) in *A. thaliana* (Pallucca et al., 2014). Four phosphorylation sites (Thr-924, Ser-931, Tyr-946, and Thr-947) in the C terminus domain of AtAHA2 are related to the interaction with 14-3-3s (Kabala & Janicka, 2023; Li et al., 2022b). For instance, phosphorylation at Thr-947 forms a binding motif (946YpTV) to interact with the 14-3-3s (Fuglsang & Palmgren, 2021), and phosphorylation at Tyr-946 and Thr-924 has been confirmed to trigger the binding between 14-3-3 and 946YpTV, thereby activating AHA2. PKS5 phosphorylates the Ser-931 of AHA2 and decreases the interaction between AHA2 and 14-3-3s (Falhof et al., 2016).

We found the interaction motifs (Thr-924/Ser-931/Tyr-946/Thr-947) between AHAs and 14-3-3s were highly conserved across in Streptophyta, mosses, and vascular plants (Fig. 2B). Previous study showed that Thr-924/Ser-931/Tyr-946/Thr-947 were not found in Rhodophyta (Falhof et al., 2016) and most Chlorophyta (Falhof et al., 2016; Zhang et al., 2019b). However, Thr-924, Tyr-946, and Thr-947 can be found in only two Chlorophyta (i.e., *Micrasterias_fimbriata* and *Zygnema sp*) out of 93 Chlorophyte algae and 30 Streptophyta through OneKP dataset (https://db.cngb.org/onekp/). The Thr-947 of AHA is essential for 14-3-3s binding in moss *P. patens* and liverwort *M. polymorpha* (Okumura et al., 2012a; Okumura et al., 2012b). The R domain of plant H^+^-ATPase may have evolved in streptophyte algae, which are important for plant terrestrialization (Steger et al., 2022). The 14-3-3s and H^+^-ATPase-coding genes were preferentially retained in the recent WGD event (Feng et al., 2021). In summary, the analysis implied that the 14-3-3-AHA pathway could be conserved in Streptophyta and land plants (Fig. 2B, C) (Falhof et al., 2016), .

#### Evolution of AHA and 14-3-3 related proteins in green plants

Genetic and molecular studies in *A. thaliana* have greatly advanced the understanding of the function of phytohormones (Waadt et al., 2022). Previous studies have revealed that AHAs and 14-3-3s are important for phytohormone pathways (e.g., AUX, ABA, and BR) (Camoni et al., 2018; Jaspert et al., 2011; Jiang et al., 2023a; Miao et al., 2022) by interacting with various members of the phytohormone signaling (Cotelle & Leonhardt, 2015) (Fig. 4). We performed a comparative analysis of phytohormone (i.e., AUX, GA, and ETH) signaling families in the 41 evolutionarily crucial green plants (Fig. S2).

**Fig. 4.**
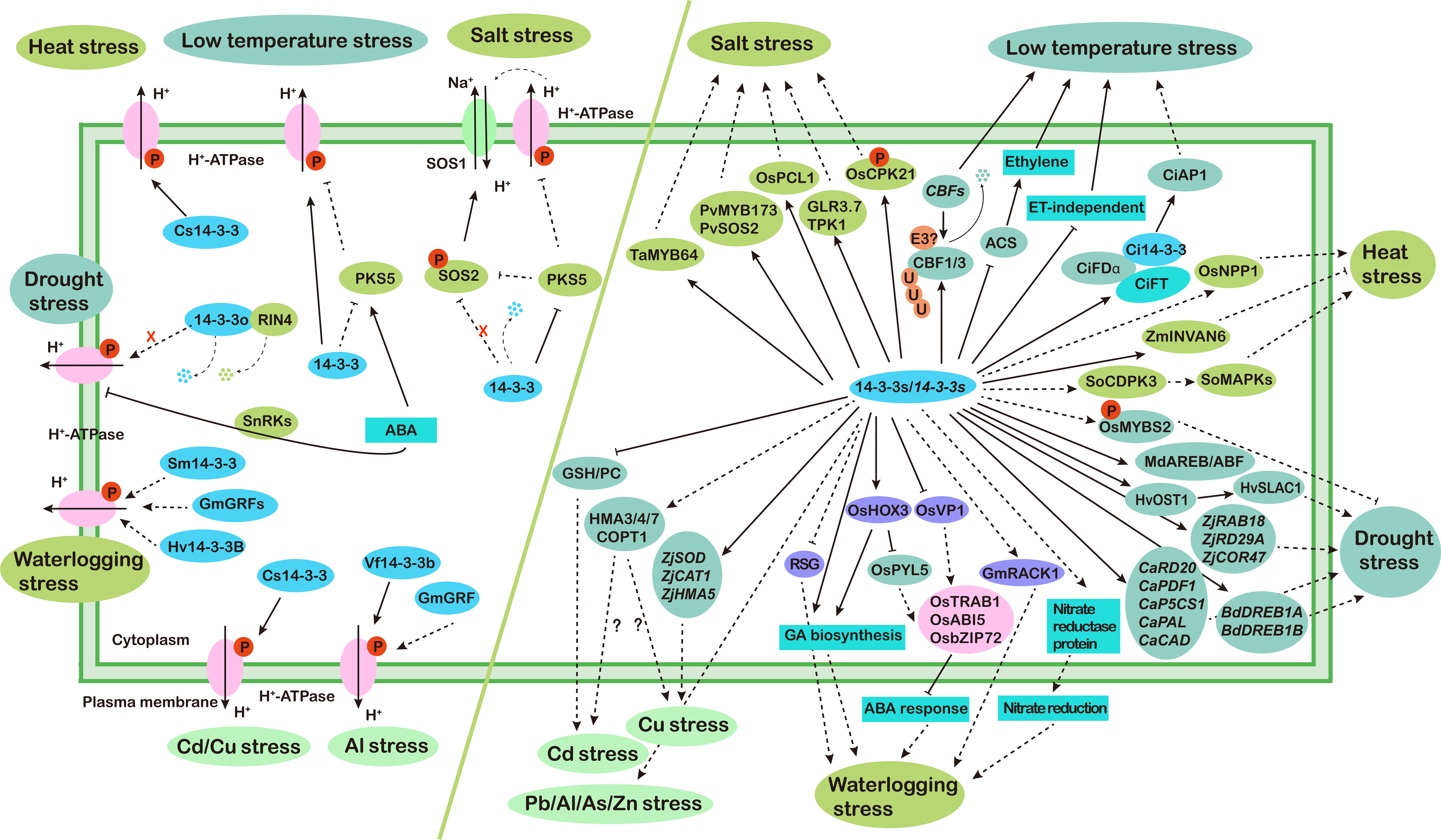
Schematic diagram of the regulation of AHA and 14-3-3 in response to key abiotic stresses in plants. Positive regulation is shown by arrows, negative regulation is indicated by lines with bars, and the dotted lines display the indirect regulation. *ZjGRF1*, seagrass *Zostera japonica 14-3-3* gene; *ZjRAB18*, *ras-related gene 18*; *ZjRD29A*, *desiccation-responsive gene 29A*; *ZjCOR47*, *cold regulated 47 gene*; MdGRF11, *Malus domestica* 14-3-3 protein; MdAREB, abscisic acid responsive elements-binding factor; MdABF, ABRE-binding factor; *BdDREB1A*, *dehydration-responsive element-binding gene 1A*; *BdDREB1B*, *dehydration-responsive element-binding gene 1B*; Hv14-3-A, *Hordeum vulgare* 14-3-3 protein; HvOST1, open stomata 1; HvSLAC1, slow anion channel 1; *Ca14-3-3B*, *Cicer arietinum 14-3-3* gene; *CaRD20*, *desiccation-responsive gene 20*; *CaPDF1*, *polypeptide deformylase 1*; *CaP5CS1*, *delta-1-pyrroline-5-carboxylate synthase 1*; *CaPAL*, *phenylalanine ammonia-lyase*; *CaCAD*, *constitutively activated cell death*; OsMYBS2, *Oryza sativa* MYB-related protein S2; RIN4, RPM1-interacting protein 4; 14-3-3o, *Arabidopsis thaliana* 14-3-3 protein; SnRKs, SNF-related serine/threonine-protein kinases. RSG, repression of shoot growth; GmRACK1, activated protein kinase C1; OsHOX3, homeobox-leucine zipper protein 3; OsVP1, viviparous 1; OsPYL5, pyrabactin resistance 1-like protein 5; OsTRAB1, bZIP transcription factor TRAB1; OsABI5, ABA-insensitive 5; OsbZIP72, basic leucine zipper 72; Hv14-3-3B, 14-3-3 protein; GmGRFs, *Glycine max* 14-3-3 proteins; Sm14-3-3, *Suaeda maritime* 14-3-3 protein. OsNPP1, nucleotide pyrophosphatase/phosphodiesterase 1; ZmINVAN6, *Zea mays* invertase alkaline neutral 6; SoCDPK3, *Spinacia oleracea* calcium-dependent protein kinase 3; SoMAPKs, Mitogen[activated protein kinases; Cs14-3-3, *Cucumis sativus* 14-3-3 protein. CiFDα, a bZIP transcription factor of citrus; CiFT, flowering locus T; CiAP1, apetala1; ACS, 1-aminocyclopropane-1-carboxylate synthetase; E3, E3 ligase; CBF, C-repeat binding factor; PKS5, SOS2-like protein kinase 5. PvSOS2, *Phaseolus vulgaris* salt overly sensitive 2; PvMYB173, MYB protein 173; TaMYB64, *Triticum aestivum* MYB protein 64; OsCPK21, calcium-dependent protein kinase 21; OsPLC1, phospholipase Cs 1; SOS1/2, salt overly sensitive 1/2; TPK1, tandem-pore K^+^ channel 1; GLR3.7, glutamate receptor 3.7. GSH, glutathione; PC, phytochelatin; HMA3/4/7, heavy metal ATPase 3/4/7; COPT1, copper transporter 1; *ZjGRF1*, seagrass *Zostera japonica 14-3-3* gene; *ZjSOD*, *superoxide dismutase*; *ZjCAT1*, *peroxidase and catalase 1*; *ZjHMA5*, *heavy metal ATPase 5*; Vf14-3-3b, *Vicia faba*14-3-3 protein; Cs14-3-3, *Cucumis sativus* 14-3-3 protein.

Using 11 AtAHAs as the reference sequences, we found 377 orthologues from 39 representative genomes of green plants (Figs. S3). Evolutionary analysis indicated that s-adenosylmethionine synthase (SAM) / (1-aminocyclopropane-1-carboxylate synthase (ACS) proteins are conserved in all green plants and seems to have originated from Rhodophyta (Fig. S2). Protein topology prediction displayed that AHA and ACS proteins were highly conserved in green plants (Fig. S4). Further investigations suggest that 24 regulatory components of phytohormone pathway had orthologues in the liverworts (e.g., *Marchantia polymorpha*) (Fig. S2). The regulatory 14-3-3 protein components and genetic similarity analysis revealed that SAMs, ethylene receptors (ETRs), GA insensitive dwarf (GIDs), auxin transporter-like proteins (LAXs), nitrilases (NITs) had the highest similarity across the green plants (Fig. S2). Most of GA signaling proteins [e.g., sleeps (SYLs) and gibberellin-regulated proteins (GASAs)] were not identified in the streptophyte algae (Fig. S2), indicating that the GA response system might have emerged from land plants (Blazquez et al., 2020). The majority of ET and IAA signaling proteins were identified in streptophyte algae (e.g., *Klebsormidium flaccidum* and *Mesotaenium endlicherianum*), suggesting that ETH and IAA signaling could align well with the appearance of land plants (Ju et al., 2015; Mutte et al., 2018). Therefore, the analysis implied that the interaction of ACS/NIT with 14-3-3s could be conserved in green plants (Jaspert et al., 2011; Yoon & Kieber, 2013).

Interaction of 14-3-3s and AHAs also play important role in plant growth and development (Fulgosi et al., 2002; Schoonheim et al., 2007). Active suppression of development and defense against stress are complementary strategies for plants to cope with distinct environments (Zhang et al., 2020). AHAs, sulfite reductase 1 (SIR1), heat shock protein (HSP), non-yellow coloring 1 (NYC1), protein-L-isoaspartate O-methyltransferases (PIMTs), galactinol synthases (GOLSs) displayed high similarity in green plants (Fig. S2). 14-3-3 proteins participate in the elongation of cells via H^+^-ATPase, which promote the excretion of protons into the cell wall (Huang et al., 2022b). Besides, the interaction between H^+^-ATPase and 14-3-3s creates a site to bind with fusicoccin (Chevalier et al., 2009), which enhance seed germination and radicle elongation through improving the uptake of K^+^ in barley (van den Wijngaard et al., 2005).

#### Comparative evolutionary and gene expression analysis of AHA and 14-3-3 genes in a eudicot, a monocot, and a moss

We analyzed the public transcriptomics data for the roles of AHAs and 14-3-3s in abiotic stresses using *A. thaliana*, barley, and *P. patens*. Barley *HvAHAs* and *HvGRFs* datasets were obtained from various stress treatments such as drought (Harb et al., 2020), submergence (Borrego-Benjumea et al., 2020), high temperature (Pacak et al., 2016), cold (Haas et al., 2020), and salinity (Fu et al., 2019). The expression of *AtAHAs* and *AtGRFs* was obtained from studies of drought (Chen et al., 2021d), low temperature (Sharma et al., 2018), heat stress (Blair et al., 2019), waterlogging (Yang et al., 2021b), and salt (Yang et al., 2018) stresses. The expression of *AHAs* and *14-3-3s* in *P. patens* was investigated via studies of drought, cold, salt (Khraiwesh et al., 2015), and heat (Elzanati et al., 2020) treatments. Raw expression data were retrieved and analyzed by fastqc (https://www.bioinformatics.babraham.ac.uk/projects/fastqc/), hisat2 (https://github.com/DaehwanKimLab/hisat2), samtools (https://github.com/samtools/samtools), and stringtie (https://ccb.jhu.edu/software/stringtie/) (Jiang et al., 2021; Pertea et al., 2016).

In *A. thaliana*, *AtAHA1/2/3* displayed high expression, while *AtAHA6/9* showed low expression under five abiotic stresses (Fig. 5A). Most *AtAHAs* were upregulated under cold stress, while downregulated under heat conditions. *AtAHA3/4* was significantly induced by salt treatments (Fig. 5A), which is in agreement with previous study that rapid regulation of the PM H^+^-ATPase activity is associated with salinity stress (Bose et al., 2015). In barley, *HvAHA2* displayed high expression, while *HvAHA6/8* showed low expression under abiotic stresses (Fig. 5A). Salinity stress decreased the H^+^-ATPase activity in root apex and mature zone of barley, and the H^+^-ATPase activity was much lower in the root apex (Shabala et al., 2016), which is in accordance to the expression of *HvAHA2* was slight decreased under salt stress (Fig. 5A). In *P. patens*, *Pp3c26_8740* displayed high expression, while *Pp3c9_14700* showed low expression under five abiotic stresses (Fig. 5A). *Pp3c23_10370* was induced by all abiotic treatments, while *Pp3c3_7160*, *Pp3c3_7180*, *Pp3c12_3320*, and *Pp3c26_8740* were reduced under all abiotic stresses, implying important roles in stresses (Khraiwesh et al., 2015). In total, we propose that that *AHAs* might be some of the direct targets under abiotic stresses.

**Fig. 5.**
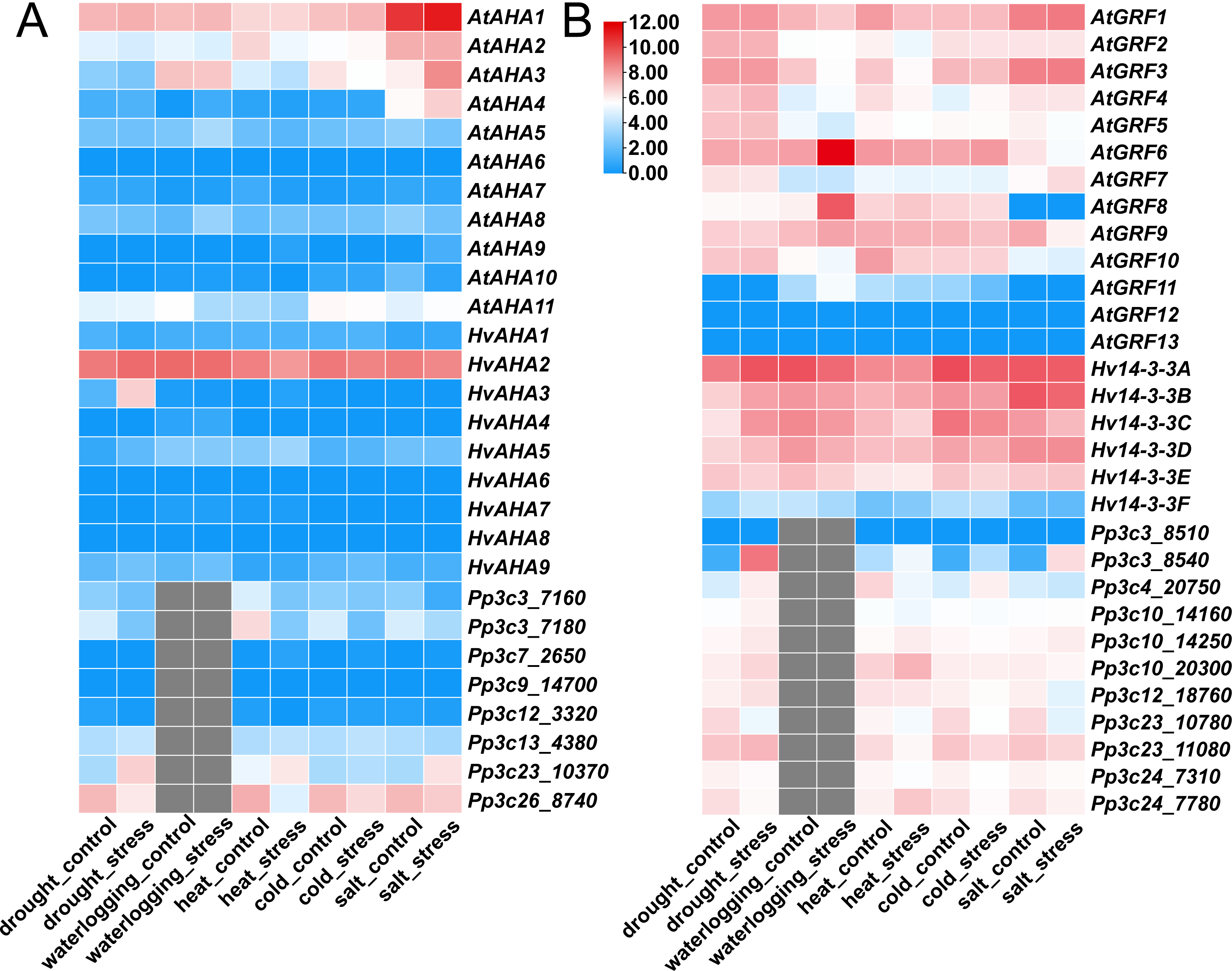
Expression of *AHA* (A) and *14-3-3* (B) genes in Arabidopsis, barley, and *P. patens* under abiotic stress. The heat map was constructed using TBtools from public databases.

*GRF12* and *GRF13* had insignificant expression under all stress, indicating that these two genes might exhibit pseudo-functionalization. *GRF8* and *GRF11* showed high expression under cold, heat, and waterlogging. In addition, *GRF4* and *GRF6* expression was found to be elevated in *A. thaliana* under low temperature stress (Fig. 5B). In addition, the transcripts of *GRF8* were upregulated by heat in *A. thaliana*. Except *Hv14-3-3F*, *Hv14-3-3s* showed high expression in drought, salt, waterlogging, cold, and heat stresses (Fig. 5B). *Hv14-3-3B* demonstrated high expression patterns under salt stress while its expression was low under heat stresses. Moreover, *Hv14-3-3s* showed low expression in heat stress compared to others signals and reduced expression under waterlogging stress (Fig. 5B).

Except *Pp3c3_8510*, other *14-3-3* genes showed high expression under salt, cold, and heat stresses in *P. patens*. Unlike *A. thaliana*, and barley, most of the *14-3-3s* in *P. patens* were upregulated during drought stress, implying their varied roles in diverse species (Jiang et al., 2023b). In addition, *Pp3c3_8540* was upregulated, while *Pp3c24_7310* and *Pp3c23_10780* were downregulated under all conditions (Fig. 5B). *Pp3c10_14250*, *Pp3c10_20300*, *Pp3c24_7780*, and *Pp3c3_8540* were induced in the heat treatments, while *Pp3c4_20750* was obviously reduced under heat stress. Moreover, except *Pp3c3_8540* and *Pp3c3_20750*, all *Pp14-3-3s* were decreased under cold stress. The protein of GRF2/5 displayed high similarity (over 85%) with Pp3c24_7780 and consistent expression pattern under low temperature and salt stress (Fig. 5B). Likewise, the high similarity (more than 82%) of orthologous was found among GRF1/3/4 and Pp3c23_11080, which illustrated a similar response to heat treatment. 10 and 13 gene pairs have shown more than 90% similarity in *A. thaliana* and *P. patens*, respectively. *Pp3c24_7310*/*Pp3c24_7780*, *Pp3c3_8510*/*Pp3c3_8540*, and *GRF2*/*GRF13* gene pairs showed tandem duplication rather than segmental duplication (Cao et al., 2016). Taken together, these results implied that there could be a crosstalk between *14-3-3s* and *AHAs* in response to diverse abiotic stresses in different plants (Figs. 5).

#### Concerted interaction of AHAs and 14-3-3 proteins in plant abiotic stresses

There are increasing evidences for these concerted actions of AHAs and 14-3-3s in response to abiotic stresses in plants (Li et al., 2022a) (Fig. 4). For instance, general control nonrepressible-4 (GCN4) could degrade RPM1-interacting protein 4 (RIN4) and GRF11 (14-3-3o), reducing the activity of H^+^-ATPase for closure of stomata that ultimately increased drought tolerance in *A. thaliana* (Kaundal et al., 2017). However, recently study showed that ABA-activated BRI1-Associated Receptor Kinase 1 (BAK1) phosphorylates at Ser-944 to activate AHA2, which can cause rapid H^+^ flux for stomatal closure (Pei et al., 2022). Under high temperature treatment, AtGRFs might modulate stomatal opening through enhancing AHA activity and phototropin (PHOT) signaling in guard cells (Kostaki et al., 2020). The 14-3-3 proteins increased activity of the H^+^-ATPase under heat stress resulted from reversible protein phosphorylation in cucumber (*Cucumis sativus* L.) (Janicka-Russak & Kabala, 2012). Moreover, long term low temperature induces the increased *AHA* gene transcription and activation of H^+^-ATPase enzymatic activity, which is due to the induction of 14-3-3 proteins (Muzi et al., 2016), and At14-3-3s regulate the H^+^-ATPase activity that protect plants against low temperature (Zhao et al., 2021). The H^+^-ATPases and 14-3-3 proteins also were molecular switch under salt stress (Feng et al., 2021). Cadmium (Cd) and copper (Cu) stress affect the activity of H^+^-ATPase through reversible protein phosphorylation, then results in the binding of 14-3-3s in *Cucumis sativus* (Janicka-Russak et al., 2012). Here, we discuss the roles of AHAs and 14-3-3s in each of the key abiotic stresses in plants.

#### The role of 14-3-3s under drought stress

The growth, development and reproduction of plants are adversely affected by drought stress (Ren et al., 2019; Wang et al., 2023b). Plant stomata controls water loss and photosynthetic CO_2_ uptake through guard cells (Chen et al., 2017b; Peng et al., 2022). Over the years, a large body of studies have proved that AHAs and 14-3-3s are crucial in guard cell signaling (Babla et al., 2019; Cotelle & Leonhardt, 2015; Li et al., 2022a; Peng et al., 2022) (Fig. S5) under drought stress (Li et al., 2022b). It is reported that 14-3-3s play a fundamental role in the drought stress in many plants including *A. thaliana*, rice, wheat, maize, barley (Figs. 4, S6). In *A. thaliana*, silencing of *AtGF14*μ (*14-3-3*μ) enhanced drought tolerance (Sun et al., 2014), and overexpression of *Glycine soja GsGF14o* in *A. thaliana* reduced drought tolerance via repressed root hair development (number and length) and stomatal size, decreasing the water uptake and transpiration rate (Liu et al., 2016; Sun et al., 2014). Moreover, PM H^+^-ATPase is a client of 14-3-3 protein interaction, and the reduction of its activity is an essential step in membrane depolarization to mediate stomatal closure in drought treatments of *A. thaliana* (Merlot et al., 2007). During stomatal closure, phosphorylation at S42 of 14-3-3 binding domain in the N-terminal controls 14-3-3 binding to tandem-pore K^+^ channel 1 (TPK1) for elevated TPK1 activity in the tonoplast in *A. thaliana* (Isner et al., 2018). Further, 14-3-3s could bind with many CPKs, which can phosphorylate both 14-3-3-binding sites in 14-3-3 clients and 14-3-3 proteins under drought stress (Ormancey et al., 2017; Yip Delormel & Boudsocq, 2019).

Rice calcium-dependent protein kinase 1 (OsCDPK1) activates one 14-3-3 protein OsGF14c to increase drought tolerance through transducing the post-germination Ca^2+^ signal (Ho et al., 2013). In addition, constitutively expressing *OsGF14f* showed drought tolerance via increasing endogenous ABA level, illustrating the pivotal roles of 14-3-3s during drought stress (Liu et al., 2016). Stress-induced F-Box protein-coding protein (OsFBX257) mediates drought stress adaptations through interacting molecularly with GF14b/c (Sharma et al., 2023). Sugar starvation-regulated protein (MYBS2) interaction with GF14b/c/d/e might be in response to drought stress in *O. sativa* (Chen et al., 2019b), and *Osgf14b* knockout mutant showed enhanced drought tolerance (Liu et al., 2019).

Recently, using firefly luciferase complementation (LUC) test and yeast two-hybrid (Y2H), we found that Hv14-3-3A interact with open stomata 1 (OST1) and slow anion channel 1 (SLAC1) in barley, which might contribute to stomatal closure in barley (Jiang et al., 2023b). Overexpression of *AtGRF6* (*14-3-3*λ) in cotton increased drought tolerance in cotton, which is due to higher photosynthesis caused by the open stomatal phenotype (Yan et al., 2004). Besides, binding of ZmGF14-6 (one 14-3-3 protein) to ZmKAT1 enhances channel open probability and modulating a large of channels in PM (Cotelle & Leonhardt, 2015; Sottocornola et al., 2008). In orange (*Citrus sinensis*), most *CsiGF14* genes were increased when exposed to drought stress (Lyu et al., 2021) and *CaGF14* of chickpea (*Cicer arietinum*) showed high expression in roots compared to shoots in response to desiccation stress (Chakraborty et al., 2022). Expression levels of *Camellia sinensis CsGRF21* were induced, while *CsGRF9* was decreased under drought stress (Zhang et al., 2022b). Altogether, these evidences might due to varied drought conditions and different strategies adopted by plants to respond to drought treatment (Huang et al., 2022b). Besides, several indications demonstrate that 14-3-3s play fundamental roles in stomatal movements during drought stress, but the precise molecular mechanisms will be identified in the future.

#### Effect of waterlogging on 14-3-3s

Flooding stress activates a range of regulatory mechanisms in plants such as modulation of chromatin structure and transcriptional activation processes (Lee & Bailey-Serres, 2021). It was demonstrated that 14-3-3s may participate in waterlogging via proteomics and gene expression (Figs. 4, S7). In *A. thaliana*, *14-3-3*κ mutants were more tolerant to hypoxia (part of waterlogging stress) relative to wild type seedlings, and 14-3-3κ negatively mediates CPK12 cytoplasm-to-nucleus translocation via decreasing ethylene signaling in hypoxia stress (Fan et al., 2023). In rice, OsGF14h can act as a signal switch to repress ABA signaling via interacting with viviparous 1 (OsVP1) and homeobox-leucine zipper protein 3 (OsHOX3) and increase GA synthesis to improve flood adaptation (Sun et al., 2022b). After flooding stress, *Glycine max* 14-3-3 (GRF7) protein expression was significantly increased in the hypocotyl of seedlings (Wang et al., 2017a) and level of 14-3-3s in PM was also altered in response to flooding in soybean (Komatsu et al., 2009). Waterlogging induced *Brassica napus Bn1433-1* at 4 h, while it was restored to normal transcript level at 8 h. In addition, the expression of *Bn1433-2/4* was decreased at 12 h (Zhan et al., 2010). 14-3-3s might inactivate repression of shoot growth (RSG) as a GA-regulated submergence escaping strategy (Bashar et al., 2019). Hypoxia activates nitrate reductase (NR) via promoting the disassociation of 14-3-3s inhibitor and NR dephosphorylation in tomato roots (Allegre et al., 2004). In *Populus tremula*, *PtGRF1/2/4a* and *12b* were strongly upregulated in leaves at 168 h of oxygen deficiency (Tian et al., 2015). The regulation of ATP synthases by 14-3-3s is a way for plant to adapt to environmental changes including anoxia in roots (Bunney et al., 2001). Nevertheless, the specific relationship between flooding stress and the 14-3-3s remain unclear.

#### The role of 14-3-3 proteins in temperature stress

The continuing increase of worldwide temperature with an average 0.3[per decade is impacting crop growth and productivity (Saini et al., 2022; Wang et al., 2017b). Heat stress responses are related to the activation of heat-shock transcription factors (HSFs) and heat-shock proteins (HSPs) in plants (Chen et al., 2021a; Shekhawat et al., 2022). Phosphorylated HspB6 (Hsp20) interact with seven isoforms of 14-3-3 protein (Muranova et al., 2022) and locking the interaction of 14-3-3s impairs heat-induced stomatal opening in *A. thaliana* (Kostaki et al., 2020). Under high temperature condition, the phosphorylation status of OsGF14s is influenced by the disruption of nucleotide pyrophosphatase/phosphodiesterase 1 (NPP1), and 14-3-3s may play a pivotal role in carbohydrate accumulation in the *npp1* mutant through interacting with metabolic enzymes (Figs. 4, S8) (Inomata et al., 2018). In addition, heat stress obviously enhances the transcripts of *OsGF14b*/c/*d* in rice (Yao et al., 2007). However, another study showed a sustained repression of *14-3-3* under high temperature stress, which modulate ATPase/synthase complex, the PM expansion, and nitrate reductase (Islam et al., 2020). Recently, we showed that that barley HvHSP90s interact with Hv14-3-3 using LUC imaging assay (Jiang et al., 2023b). In maize, cytosolic invertase INVAN6 interacts with seven 14-3-3 proteins (Zm14-3-3_b/c/f/i/m/q/z) to play a fundamental role in the meiotic progression during heat stress (Huang et al., 2022a). In addition, the application of cyclic adenosine monophosphate increased the expression of 14-3-3-like protein GF14-6/12 under heat stress in maize (Liang et al., 2022; Yang et al., 2021a). In spinach (*Spinacia oleracea* L.), heat stress-responsive proteins (e.g., CDPK, MAPK) were reported to function in 14-3-3-mediated signaling pathway (Zhao et al., 2018). Moreover, it was shown that 14-3-3s have positive or negative effects on the heat tolerance of plants in the same genus. For instance, 14-3-3 protein was significantly upregulated in *Lilium longiflorum* while down-regulated in *Lilium distichum* under high temperature stress (Fu et al., 2020). In wheat, the down-regulation of 14-3-3 protein was observed after heat stress, indicating that 14-3-3 negatively regulates stress response (Laino et al., 2010; Yang et al., 2011). Taken together, it would be useful to explore the function of the 14-3-3s and their client proteins under heat stress from transcriptional and post-translational modifications (PTM).

Low temperature stress is an acute threat that limits plant growth (Ding et al., 2020; Jiang et al., 2020). The research work on 14-3-3s in response to low temperature is well certified and mainly focused on *A. thaliana* (Figs. 4, S9). Overexpression of *At14-3-3*ε and *At14-3-3*ω enhance low temperature tolerance, and H^+^-ATPase activity was induced in 14-3-3ω-overexpressing plants (Visconti et al., 2019). Cold-responsive protein kinase 1 (CRPK1) phosphorylates 14-3-3s (Guan et al., 2023) and 14-3-3 proteins are shuttled from cytosol to nucleus to regulate C-repeat binding factor (CBF) transcription factor under cold stress in *A. thaliana* (Liu et al., 2017). Besides, BYPASS1-LIKE (B1L) enhances low temperature tolerance of *A. thaliana* possibly through 14-3-3λ and stabilization of CBF3 (Chen et al., 2019a). In *A. thaliana*, *RARE COLD INDUCIBLE 1A* (*RCI1A*/*GRF3/14-3-3*ψ) and RCI1B are regulators of cold acclimation and constitutive freezing tolerance (Catala et al., 2014) and the transcription level of *RCI1A* and *RCI1B* were higher in cold stress but lower under other abiotic stresses, implying a specific function for their response to cold (Abarca et al., 1999). *A. thaliana* BRI1-Flag (*14-3-3*ψφε) (overexpressing Flag-tagged BRI1 in *14-3-3*ψφε knockout mutant) displayed significantly reduced cold tolerance when compared to wild type (Lee et al., 2020). In rice, the RING E3 Ub ligase OsATL38 negatively modulates cold stress through mono-ubiquitination of OsGF14d, while OsGF14d positively regulates the low temperature stress (Cui et al., 2022). OsGF14h interacts with mother of FT and TFL 2 (MFT2) and bZIP-type transcription factor (OREB1) under temperature conditions via modulating ABA-responsive genes (Yoshida et al., 2022). The cold stress-induced upregulation of *14-3-3s* includes *CaGF14f* in *Cicer arietinum* (Chakraborty et al., 2022), Hv14-3-3s in *H. vulgare* (Golebiowska-Pikania et al., 2017; Wojcik-Jagla et al., 2020), *VviGRFs* (*VviGRF9b/14/15/17/18/like2*) in grape (Cheng et al., 2018) and *AhGF14* in *Arachis hypogaea* (Chakraborty et al., 2022). In *Citrus*, a bZIP transcription factor CiFDα responds low temperature through forming a florigen activation complex with Ci14-3-3 and flowering locus T (FT) (Ye et al., 2022). 14-3-3 proteins could bind to cold-induced cytosolic Ca^2+^ to modulate PKS5 activity that might help decode Ca^2+^ signaling in the response to cold stress in soybean (Sun et al., 2022a). Most interestingly, the interaction of AHAs and 14-3-3s for cold tolerance in plants were also clearly reported. In maize roots, 14-3-3 proteins were involved in low-temperature responses by binding to AHA to activate the proton pump which causes K^+^ influx and water uptake (Cao et al., 2016). In pepper (*Capsicum annuum*), 14-3-3-like protein D/E and 14-3-3 protein 7 were significantly down-regulated by the application of 24-epibrassinolide (EBR), implying their role in EBR-induced growth and possible inhibitory interaction between PM H^+^-ATPase and the dimer of 14-3-3 protein under chilling stress (Li et al., 2021). Together, the 14-3-3 proteins could respond to low temperature stress via different regulatory components (e.g., phytohormone signaling pathway, H^+^-ATPase, and Ca^2+^) (Huang et al., 2022b).

#### Differential responses of 14-3-3s to salt stress

Soil salinity is one of the limiting factors that strongly affects crop quality and growth world-wide (Chen et al., 2021b; Zhou et al., 2014). In recent years, many studies have exhibited the response of 14-3-3s to salt stress (Figs. 4, S10). The *A. thaliana* 14-3-3λ and 14-3-3κ have been identified to negatively modulate salt tolerance via interacting with SOS2. In control, two 14-3-3 proteins interacted with and repressed SOS2 activity to repress the SOS pathway, while salt stress decreased the interaction between SOS2 and 14-3-3s (Zhou et al., 2014). Furthermore, salt condition induces the degradation of 14-3-3λ/κ, partially via the actions of SOS3-like calcium binding protein 8 / calcineurin-B-like10 (SCaBP8/CBL10), and decreases the PM localization of 14-3-3λ (Tan et al., 2016). Furthermore, salt stress enhances the interaction between PKS5 and 14-3-3s, repressing its activity and alleviating inhibition of the SOS2 on 14-3-3s (Yang et al., 2019). Under salt treatment, overexpressing SOS1 C-term in *A. thaliana* plants displayed more salt tolerance via sequestration of inhibitory 14-3-3s, thus decreasing Na^+^ accumulation in leaves, proline, soluble sugar, and starch levels to increase biomass (Duscha et al., 2020). Salinity treatment also increases 14-3-3 phosphorylation by Ca^2+^-dependent protein kinases 3 (CPK3), thereby contributing to an additional regulation of tonoplast TPK1 channels in *A. thaliana* (Latz et al., 2013). 14-3-3s may modulate the GORK activity via activation and binding with CPK3/6/21/23 particularly CPK21 under salinity stress (van Kleeff et al., 2018). Glutamate receptor-like protein 3.7 (GLR3.7) might interact with14-3-3ω proteins in response to salt stress via calcium signaling (Wang et al., 2019).

In rice, OsCPK21, a calcium-dependent protein kinase, interacts with OsGF14e to improve plants’ response to salt stress via phosphorylating at Tyr-138 of 14-3-3 protein (Chen et al., 2017a). OsGF14b positively mediates rice salt tolerance through interacting with phospholipase C1 (OsPLC1) and inhibiting the ubiquitination and hydrolysis of OsPLC1 to increase its activity and stability under NaCl stress (Wang et al., 2023a). Besides, the roles of 14-3-3s in plant salt stress were identified by the salt-induced upregulation of *TOMATO 14-3-3 PROTEIN 1/4/7/10* (*TFT1/4/7/10*) (Xu & Shi, 2006), *AhGF14e/g/h/j/l* in *Arachis hypogaea* (Chakraborty et al., 2022), and the interaction between PvMYB173/PvSOS2 and PvGF14a/g in *Phaseolus vulgaris* (Li et al., 2018). However, overexpression of apple *MdGRF6* in apple calli and tobacco decreased tolerance to salt stress, which might be associated with salt stress-related genes (e.g., *MdSOS2/3*, *MdCBL-1*, *MdWRKY30*, and *MdMYB46*) (Zhu et al., 2023). In summary, 14-3-3s influence plants salt stress potentially through the SOS signaling pathway, ion transport, and the ROS-scavenging system (Huang et al., 2022b). It will be highly valuable to validate the molecular function of client proteins of 14-3-3s in salt stress response in different plant species.

#### 14-3-3s participate in heavy metal stress response and detoxification

Heavy metals and metalloids such as Cd, Cu, aluminium (Al), zinc (Zn), arsenic (As), and lead (Pb) have serious effects on plants when their concentrations are too high in soil and water (Deng et al., 2021; Tang et al., 2023). Plant 14-3-3 proteins also participate widely in the response to heavy metals (Figs. 4, S11). In *A. thaliana*, a putative binding motif of 14-3-3 was confirmed in the C-terminal cytosolic domain of cadmium/zinc-transporting ATPase 3 (HMA3) *in silico* (Kramer et al., 2007). It was reported that heavy metal ATPase 4 (HMA4) may interact with 14-3-3s through tandem affinity purification (TAP) of 14-3-3 protein complexes (Chang et al., 2009). Besides, 14-3-3 affinity chromatography (AC) of plant extracts showed it could interact with HMA7 (heavy metal ATPase 7, copper-transporting ATPase) (Shin et al., 2011). Besides, yeast two-hybrid demonstrated that copper transporter 1 (COPT1) was characterized as putative 14-3-3 interactors in *A. thaliana*. It has been reported that *At14-3-3*κ*/*λ negatively modulate plant Cd stress via mediating the stimulation of glutathione and ROS (Zhang et al., 2019a). The *14-3-3*κλ double mutant demonstrates greater tolerance to Cd through increasing the content of phytochelatin (PC) and glutathione (GSH), and overexpression of 14-3-3κ or 14-3-3λ decrease Cd tolerance (Zhang et al., 2019a). Furthermore, overexpression of *Zostera japonica ZjGRF1* increases copper tolerance in transgenic *A. thaliana* through enhancing *HMA5*, *SOD*, and *CAT1* expression levels (Chen & Qiu, 2022).

For other plant species, the transcripts of *OsGF14c/e/g* have significantly reduced at 1 day of As condition, whereas after 3 days of As treatment, maximum downregulation (2.8-fold) of expression was displayed in *OsGF14a*, followed by *OsGF14b/c/d/h* in rice (Pathare et al., 2016). In barley, the nitric oxide (NO) donor sodium nitroprusside reduced Cd-modulated seedling growth inhibition through increasing the relative abundance of Hv14-3-3 protein (Alp et al., 2022). The transcript abundance of *14-3-3* was upregulated in Zn treatment in tea (*Camellia sinensis* L.) root and leaf (Mukhopadhyay et al., 2013). Highly increased levels of 14-3-3 protein were observed at 4 h and 8 h of Cu stress in *Phanerochaete chrysosporium* (Okay et al., 2020). In addition, proteomics data indicated that 14-3-3 proteins might participate in Pb stress mechanisms in *Paspalum fasciculatum* (Salas-Moreno et al., 2022a). Micromolar magnesium could mitigate the toxicity of Al through promoting the phosphorylation of VHA2 (PM H^+^-ATPase) and improving the interaction between VHA2 and 14-3-3b in *Vicia faba*. L (Chen et al., 2015). *Coprinus atramentarius* can tolerate and accumulate Cd by stimulating the content of 14-3-3s and activating the expression of *14-3-3s* (Xie et al., 2017). Proteome and phosphoproteome analysis showed that exogenous nitrogen enhances Cd tolerance partially through accumulating 14-3-3 proteins in poplar (*Populus yunnanensis*. L) plants (Huang et al., 2019). Abundances of ATPases and 14-3-3-like protein were obviously modified under Pb treatment in poplar roots, and Pb uptake and sequestration might be associated with the interaction between H^+^-ATPase and 14-3-3 protein in poplar (Szuba et al., 2020). Three 14-3-3-like protein (e.g., GF14-B and GF14-D) were upregulated in response to Cd treatment in *P. fasciculatum* leaves (Salas-Moreno et al., 2022b). However, the precise molecular mechanisms of 14-3-3s in heavy metal toxicity remain elusive.

#### Integrated function of 14-3-3 proteins for plant tolerance to multiple abiotic stresses

Increasing evidences demonstrated that 14-3-3s can respond to diverse abiotic stresses (Huang et al., 2022b) (Figs. 4, 5). For instance, transcripts of *OsGF14g/f* were significantly increased by drought, salt, and cold stress (Yashvardhini et al., 2018). The expression of the *OsGF14b/c/e* in roots is repressed in the initial phase of stress and then induced at 24 h in drought and salt stresses (Huang et al., 2022b). Furthermore, overexpression of the wheat *TaGF14b* in tobacco confers salt/drought tolerance by regulating ABA signaling pathway (Zhang et al., 2018). Overexpression of apple *MdGRF11* (Ren et al., 2019) and *MdGRF13* (Ren et al., 2023) in *A. thaliana* display enhanced tolerance to salt and drought stress, which could due to stress-responsive proteins and antioxidant enzyme activities. Interestingly, the interaction between 14-3-3 and CPK is participated to salt/heat stresses (Fig. 6), indicating that this pathway might share in distinct abiotic stresses. OST1/14-3-3 could mediate CBF in response to cold stress (Zhang et al., 2022a) and OST1 interacts with 14-3-3 might participate to drought stress in barley (Jiang et al., 2023b). Therefore, many *14-3-3s* can respond to diverse stresses, and overexpression of these *14-3-3s* usually confer tolerance to multiple abiotic stresses. It would be interesting to investigate whether these *14-3-3s* could integrate distinct stress signaling pathways for improving crop tolerance to multiple abiotic stresses in the future.

## Conclusions and future perspectives

14-3-3s and AHAs are two essential protein families that attract significant research interests in plant molecular biology and physiology for many years. Both 14-3-3s and H^+^-ATPases are targets of distinct protein kinases and other interacting partners of plant development and environmental cues. A large of advances have been made to improve the understanding of regulatory mechanisms underlying the important functions of 14-3-3s and AHAs in response to plant abiotic stresses. We proposed that the crucial components of 14-3-3s and AHA regulatory patterns and transduction pathways are highly evolutionarily conserved in Bryophyta, Pteridophyta and Spermatophyta (Figs. 1, 2, 3, S2). Moreover, the 14-3-3-AHA pathway may have evolved from Streptophyta and undergone expansion in the Bryophyta such as mosses and liverworts (Figs. 2, 3). Besides, 14-3-3s and AHAs can function as hubs of diverse signaling pathways to integrate different phytohormone signals in plant cell responding to abiotic stresses (Figs. 4, 5, S2). In addition, there could be coordination and crosstalk between 14-3-3s and H^+^-ATPases in plant abiotic stress responses (Figs. 4, 5).

Future research work should focus on the molecular evolution and functions of 14-3-3s, AHAs, and their respective potential clients. It will be of importance to investigate their functional redundancy versus isoform specificity, and to reveal the environmental stimuli (e.g., waterlogging and heat stress) that impact these processes. Also, protein dephosphorylation and phosphorylation are the most essential regulatory mechanisms for 14-3-3s, AHAs and other interacting partners (client proteins). Further investigations are needed to understand the specificity of phosphatases and protein kinases that target the phosphosites of 14-3-3s and H^+^-ATPases, especially at the C-terminal. Moreover, technological advances in cryogenic electron microscopy (cryo-EM), multi-isotope imaging mass spectrometry (MIMS), single-particle analysis, and X-ray free electron laser (XFEL) will improve investigation the role of the 14-3-3-client protein complexes through high-resolution crystal structure. This is useful in exploring the dynamic changes of 14-3-3s subcellular location and identifying new binding motifs between H^+^-ATPases and 14-3-3s. Finally, investigation the functional and regulatory mechanisms of 14-3-3s and H^+^-ATPases needs to be determined by genome editing and genetic engineering for improving abiotic stress tolerance in crops and for exploring the burning questions in evolution and ecology of evolutionarily important early divergent model plants and streptophyte algae. Taken together, these approaches will elucidate the comprehension of the molecular evolution and function of 14-3-3s and H^+^-ATPases in adapting to the different abiotic stresses in land plants.

## Acknowledgements

We thank Zhenghong Huang (Yangtze University) and Wen Li (Yangtze University) for their contribution in data collection from public data sourice.

## Author contributions

FD and Z-HC conceived the study. WJ analyzed the data and prepared all the Figures together with GC and TT. WJ, FZ, and Z-HC analyzed the results and wrote the manuscript with support from JH, MB, TW, TT, AR, FZ, YQ, and GC. WJ, FD, and Z-HC conducted the final editing of the manuscript. All authors read and approved the final version of the manuscript.

## Data availability

All data supporting the findings of this study are available within the paper and within its supplementary materials published online.

## Supplementary materials

**Fig. S1.** Motif alignment of 14-3-3 proteins in moss, fern, rice, barley, and Arabidopsis.

**Fig. S2.** Similarity heat map AHA and 14-3-3 related proteins in spermatophyte, pteridophyte, bryophyta, and algae.

**Fig. S3.** Number of plant growth and development and hormone signaling associated with 14-3-3 proteins in different plant and algal species.

**Fig. S4.** Predicted 3D structure of AHA and ACS proteins in land plants and algal species.

**Fig. S5.** Schematic diagram of the regulation of 14-3-3 and AHA in guard cell signaling.

**Fig. S6.** Schematic diagram of the regulation of 14-3-3 in response to drought stress in plant.

**Fig. S7.** Schematic diagram of the regulation of 14-3-3 in response to waterlogging stress in plant.

**Fig. S8.** Schematic diagram of the regulation of 14-3-3 in response to heat stress in plant.

**Fig. S9.** Schematic diagram of the regulation of 14-3-3 in response to low temperature stress in plant.

**Fig. S10.** Schematic diagram of the regulation of 14-3-3 in response to salt stress in plant.

**Fig. S11.** Schematic diagram of the regulation of 14-3-3 in response to heavy metal stress in plant.

